# Lis1 promotes the formation of maximally activated cytoplasmic dynein-1 complexes

**DOI:** 10.1101/683052

**Authors:** Zaw Min Htet, John P. Gillies, Richard W. Baker, Andres E. Leschziner, Morgan E. DeSantis, Samara L. Reck-Peterson

## Abstract

Cytoplasmic dynein-1 is a molecular motor that drives nearly all minus-end-directed microtubule-based transport in human cells, performing functions ranging from retrograde axonal transport to mitotic spindle assembly^1,2^. Activated dynein complexes consist of one or two dynein dimers, the dynactin complex, and an “activating adaptor”, with maximal velocity seen with two dimers present (Fig. 1a)^3-6^. Little is known about how this massive ∼4MDa complex is assembled. Using purified recombinant human proteins, we uncovered a novel role for the dynein-binding protein, Lis1, in the formation of fully activated dynein complexes containing two dynein dimers. Lis1 is required for maximal velocity of complexes activated by proteins representing three different families of activating adaptors: BicD2, Hook3, and Ninl. Once activated dynein complexes have formed, they do not require the presence of Lis1 for sustained maximal velocity. Using cryo-electron microscopy we show that human Lis1 binds to dynein at two sites on dynein’s motor domain, similar to yeast dynein^7^. We propose that the ability of Lis1 to bind at these sites may function in multiple stages of assembling the motile human dynein/ dynactin/ activating adaptor complex.

---

Cytoplasmic dynein-1 (dynein) is responsible for the long-distance transport of nearly all cargos that move towards the minus ends of microtubules, including organelles, membrane vesicles, and protein and RNA complexes in many eukaryotic cells. Dynein also has roles in cell division, and viruses hijack dynein for their transport^2^. Mutations in components of the dynein machinery cause neurodevelopmental and neurodegenerative diseases^8^. Activated human dynein is a large ∼4MDa multi-subunit complex composed of one or two dynein dimers (each dynein dimer contains two motor subunits and two copies each of five additional subunits), the dynactin complex (composed of 23 polypeptides) and one of several classes of dimeric, coiled-coil-containing activating adaptors (Fig. 1a)^2-6,9^. The dynein motor-containing subunit, or heavy chain, is an ATPase containing six AAA+ domains and a microtubule-binding domain that emerges from a long coiled-coil “stalk” (Fig. 1b).

**Figure 1.**
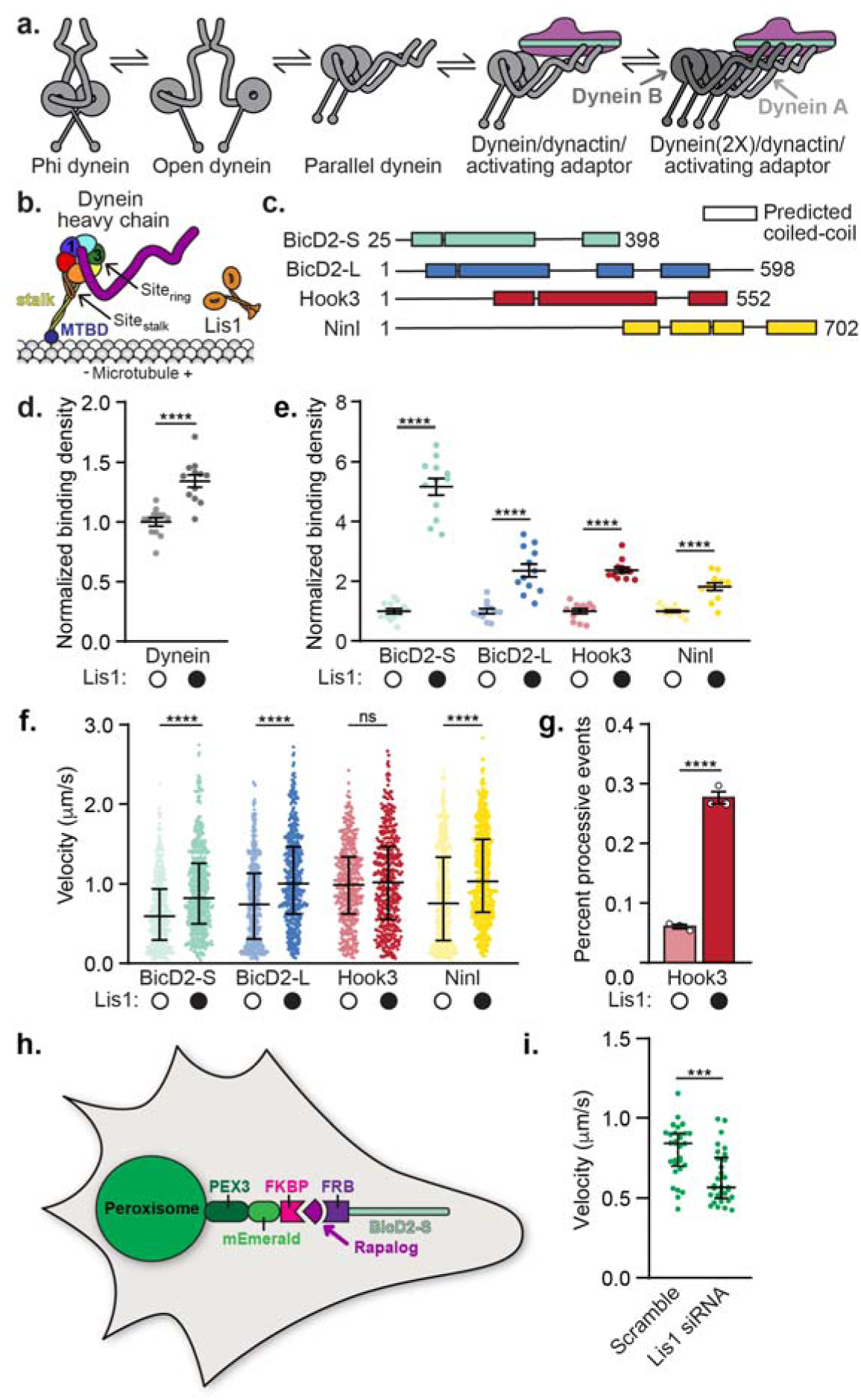
Lis1 increases microtubule binding and the velocity of activated dynein complexes. **a**. Schematic of the current model for dynein activation. Dynein is autoinihibited in the Phi conformation, opens and then adopts a parallel conformation that is seen in the activated dynein complex, which includes dynactin and an activating adaptor. Maximally activated dynein contains two dynein dimers (dynein A and B, far right) **b**. Schematic of the dynein motor domain with AAA+ ATPase domains colored in rainbow, highlighting the two Lis1 binding sites in yeast dynein, “site_ring”_ at AAA3/4 and “site_stalk_” on dynein’s stalk, which leads to dynein’s microtubule binding domain (MTBD). Lis1 is shown in orange to the right **c**. Schematic of the activating adaptor constructs used in this study. **d**. Binding density (mean ± s.e.m. dynein) of dynein alone on microtubules in the absence (white circles) or presence (black circles) of 300 nM Lis1. Data was normalized to a density of 1.0 in the absence of Lis1. Statistical analysis was performed using a two-tailed unpaired t test; ****, p<0.0001; n = 12 replicates for each condition. **e**. Binding density (mean ± s.e.m) of dynein/ dynactin/ activating adaptor complexes on microtubules in the absence (white circles) or presence (black circles) of 300 nM Lis1. The activating adaptors used are indicated. Data was normalized to a density of 1.0 in the absence of Lis1. Statistical analysis was performed using a two-tailed unpaired t test; ****, p<0.0001; n = 12 replicates for each condition. **f**. Velocity of dynein/ dynactin/ activating adaptor complexes in the absence (white circles) or presence (black circles) of 300 nM Lis1. The median and interquartile range are shown and the activating adaptors used are indicated. Statistical analysis was performed using a two-tailed Mann-Whitney test; ****, p<0.0001; ns, p=0.3498; n (individual single molecule events) = 506 (BicD2-S no Lis1), 569 (BicD2-S with Lis1), 496 (BicD2-L no Lis1), 505 (BicD2-L with Lis1), 454 (Hook3 no Lis1), 471 (Hook3 with Lis1), 490 (Ninl no Lis1), 582 (Ninl with Lis1). **g**. Percent processive runs (mean ± s.e.m.) of dynein/ dynactin/ Hook3 complexes in a higher salt buffer (60 mM KOAc versus 30 mM KOAc in our standard motility buffer) in the absence (white circles) or presence (black circles) of 300 nM Lis1. Statistical analysis was performed using a one-way ANOVA and Tukey’s multiple comparisons test; ****, p<0.0001; n = 3 replicates per condition. **h**. Schematic of the peroxisome relocation assay. **i**. Peroxisome velocity in human U2OS cells with scrambled or Lis1 siRNA knockdown. The median and interquartile range are shown. Statistical analysis was performed using a two-tailed unpaired t test; ***, p=0.0002; n (average peroxisome velocity per cell) = 30 per condition. More than 7 events were measured per cell.

Mammalian dynein (but not yeast dynein^10^) is largely immotile in the absence of dynactin and an activating adaptor^3,4,11^. Activating adaptors also link dynein/ dynactin either directly or indirectly to its cargos^2,9^. Nearly a dozen activating adaptors have been described; they share little sequence identity, but all contain a long stretch of coiled-coil that spans the ∼40 nm length of dynactin^2,9^. All activated dynein complexes that have been investigated structurally can bind two dynein dimers^5,6^, referred to as “dynein A” and “dynein B” (Fig. 1a).

Mammalian dynein in the absence of these other components adopts a conformation known as “Phi”^12,13^. Phi dynein is autoinhibited and cannot interact with microtubules productively^12^. The current model for dynein activation proposes that Phi dynein must first adopt an “Open” conformation and then ultimately a “Parallel” conformation that is observed when it is bound to dynactin and an activating adaptor (Fig. 1a)^5,6^. Little is known about how dynein switches between the autoinhibited Phi conformation and the Open and Parallel conformations that lead to the assembly of the motile activated dynein/ dynactin complex.

Genetic studies in model organisms place the dynein-binding protein, Lis1, in the dynein pathway^14-16^. Lis1 is required for many dynein functions ranging from organelle trafficking (for example^17-20^) to nuclear migration/positioning (for example^15,21-23^) to RNA localization (for example^24^). The *LIS1* gene is mutated in the neurodevelopmental disease type-1 lissencephaly^25^, and was first directly linked to dynein through genetic studies in the filamentous fungus *Aspergillus nidulans*^15^. Lis1 is a dimer of β-propellers^26,27^ and yeast Lis1 binds dynein at two distinct sites in the dynein motor domain^7,28,29^. One site is located on dynein’s ATPase ring near AAA3 and AAA4 (“site_ring_”) and the other is located on dynein’s stalk (“site_stalk_”), which leads to its microtubule-binding domain^7,28,29^(Fig. 1b.). In yeast, binding of Lis1 to dynein at site_ring_ causes tight microtubule binding and decreased velocity^7,29^, whereas binding at both sites leads to weak microtubule binding and increased velocity^7^. Lis1 also increases the binding of mammalian dynein to microtubules^30,31^ and increases the velocity of mammalian dynein/ dynactin complexes containing the BicD2 activating adaptor^32,33^. How Lis1 exerts these effects on mammalian dynein is unknown. It is also unknown if Lis1 has the same effects on dynein/ dynactin bound to other activating adaptors.

To determine how Lis1 regulates activated human dynein complexes we purified human dynein^3^ and Lis1 from insect cells, dynactin from human HEK-293T cells^34^, and the human activating adaptors BicD2, Hook3, and Ninl from *E. coli* (Supplementary Fig. 1a). Since some activating adaptors are known to be autoinhibited^35^, we chose to use well-characterized carboxy-terminal truncations of BicD2, Hook3 and Ninl^3,4,34^ (Fig. 1c). For BicD2 we made both short (BicD2-S) and long (BicD2-L) forms, which have been shown to activate dynein in vitro^3,4^ and in cells^18,36^, respectively.

We first determined the effects of Lis1 on the microtubule binding properties of dynein alone and dynein/ dynactin bound to different activating adaptors using a single-molecule assay^7^. Lis1 increased the microtubule binding density of dynein alone (Fig. 1d), consistent with studies of yeast^7,29^ and mammalian^30,31^ dynein. Lis1 also increased the microtubule binding density of dynein/ dynactin complexes bound by the activating adaptors BicD2-S, BicD2-L, Hook3, and Ninl (Fig. 1e).

Next, we determined the effect of Lis1 on the velocity of activated dynein complexes. In agreement with some previous studies^32,33^, we found that Lis1 increased the velocity of dynein/ dynactin / BicD2-S complexes (Fig. 1f). We also found that Lis1 increased the velocity of dynein/ dynactin activated by BicD2-L and Ninl, but not Hook3 in our standard motility assay buffer (Fig. 1f). However, when we raised the salt concentration in our assay buffer we found that Lis1 increased the percentage of dynein/ dynactin/ Hook3 microtubule binding events that are processive (Fig. 1g) and also increased the velocity of these processive runs (Supplementary Fig. 1b). We interpret this difference in sensitivity to the ionic strength of our assay conditions as an indication that Hook3 may have a higher affinity for dynein B compared to BicD2 and Ninl. Overall this data shows that Lis1 increases both microtubule binding and motility of dynein/ dynactin complexes bound by activating adaptors from three different families.

We next asked if Lis1 had a similar effect on activated dynein velocity in cells using a well-established peroxisome relocation assay (Fig. 1h)^36,37^. We co-transfected human U2OS cells with two constructs: 1) the rapamycin binding protein FRB fused to BicD2-S and 2) the rapamycin binding protein FKBP fused to both mEmerald and the peroxisome targeting protein Pex3 (Fig. 1h). In U2OS cells peroxisomes rarely move, but upon the addition of rapalog, which causes FRB and FKBP to interact we observed many processive runs. This is an indication that BicD2-S recruits and activates dynein/ dynactin (Supplementary videos 1-4)^37^. We quantified the velocity of these runs in cells with or without Lis1 knockdown by siRNA (Supplementary Fig. 1c) and observed a significant decrease in peroxisome velocity in Lis1 knockdown cells (Fig. 1i). These results suggest that in a cellular environment the presence of Lis1 also increases the velocity of activated dynein complexes.

We next sought to determine where Lis1 binds the human dynein motor. Experiments with the yeast homologs of dynein and Lis1 have shown that Lis1 binds dynein at two sites in the dynein motor domain (on the ring at AAA3/4 and on the stalk; Fig. 1b)^7,28,29^, although previous studies with mammalian proteins reported interactions with other regions of dynein^38,39^. Here we used cryo-electron microscopy (cryo-EM) to identify the Lis1 binding sites on human dynein. We purified monomeric human dynein motor domains and mixed them with dimeric human Lis1 in the presence of ATP-vanadate.

Although the strong preferred orientation adopted by the sample prevented us from obtaining a three-dimensional reconstruction, we generated two-dimensional (2D) class averages of the dynein/ Lis1 complex; these averages showed high-resolution features in both dynein and Lis1 (Fig. 2a). To determine whether the binding sites for Lis1 are similar in both human and yeast dynein, we compared our experimental class averages with calculated 2D projections of a model of human dynein bound to Lis1. To make this model we combined the structure of human dynein-2 bound to ATP-vanadate (PDB: 4RH7^40^) with a homology model of human Lis1 bound to dynein at the two binding sites observed with the yeast proteins (PDB: 5VLJ^7^) (Fig. 2b). To highlight the densities corresponding to Lis1, we also calculated 2D projections of human dynein-2 alone in the same orientations as our experimental data (Fig. 2c). The correspondence between our data and the model with two Lis1s bound (Fig. 2a,b) suggests that the yeast and human Lis1 binding sites are in similar regions of the dynein motor domain on the ring at AAA3/4 and on the stalk, although in the absence of a high-resolution 3D reconstruction we cannot map the exact locations on either dynein or Lis1.

**Fig. 2.**
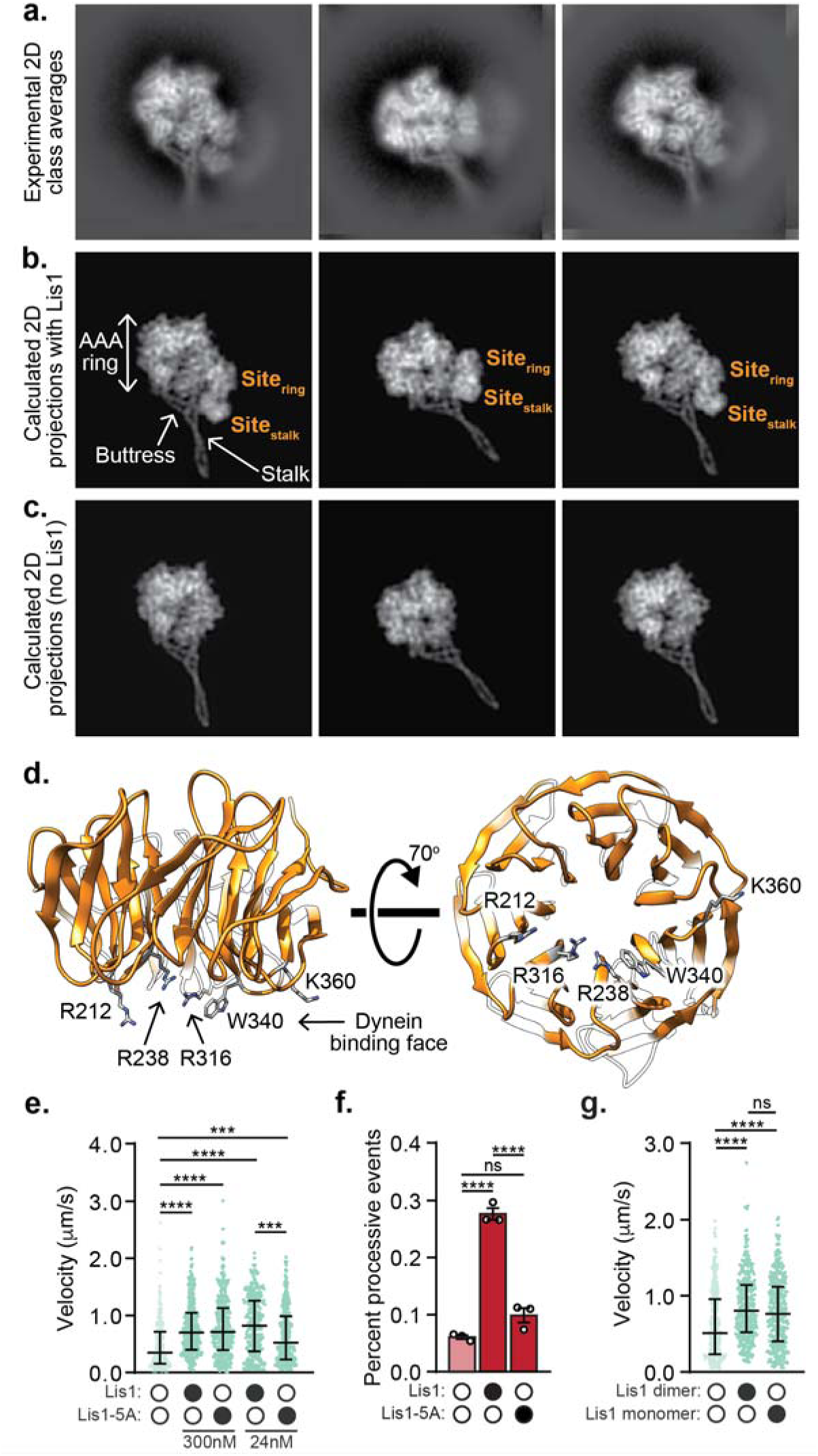
Human Lis1 binds the human dynein motor at AAA3/4 and the stalk. **a**. 2D class averages of the human dynein/ Lis1 complex in the presence of ATP-vanadate. **b**. Best-matching projections of a model combining human dynein-2 bound to ATP-vanadate (PDB: 4RH7) with homology models of human Lis1 at the locations where Lis1 binds to yeast dynein (AAA3-Walker B) in the presence of ATP-vanadate (PDB: 5VLJ). The two Lis1’s, at the two sites (“site_ring_” and “site_stalk_”) identified in yeast dynein are labeled, as are dynein’s AAA ring, stalk and buttress. **c**. Projections of 4RH7 alone in the same orientations as those shown in (b) for the model. **d**. Homology model of human Lis1 (from SWISS-MODEL) showing the five residues mutated to alanine in “Lis1-5A”. **e**. Velocity of dynein/ dynactin/ BicD2-S complexes in the absence (white circles) or presence (black circles) of Lis1 or Lis1-5A. The median and interquartile range is shown and the concentrations of Lis1 or Lis1-5A used are indicated. Statistical analysis was performed using a Kruskal-Wallis test with Dunn’s multiple comparisons test; ****, p<0.0001; ***, p=0.0002; n (individual single molecule events) = 278 (no Lis1), 344 (300 nM WT Lis1), 306 (300 nM Lis1-5A), 350 (24 nM WT Lis1), 331 (24 nM Lis1-5A). **f**. Percent processive runs (mean ± s.e.m.) of dynein/ dynactin/ Hook3 complexes in a higher salt buffer in the absence (white circles) or presence (black circles) of Lis1 or Lis1-5A. Data in the presence and absence of WT Lis1 was also presented in Fig. 1g. Statistical analysis was performed using a one-way ANOVA and Tukey’s multiple comparisons test; ****, p<0.0001; ns, p=0.0688; n = 3 replicates per condition, percentages were derived from >100 single molecule events per replicate. **g**. Velocity of dynein/ dynactin/ BicD2-S complexes in the absence (white circles) or presence (black circles) of 300 nM Lis1 dimer or 600 nM Lis1 monomer. 300 nM Lis1 dimer is equivalent to 600 nM Lis1 monomer in terms of the concentration of □-propellers. The median and interquartile range are shown. Statistical analysis was performed using a Kruskal-Wallis test with Dunn’s multiple comparisons test; ****, p<0.0001; ns, p=0.0906; n (individual single molecule events) = 364 (no Lis1), 377 (Lis1 dimer), 348 (Lis1 monomer).

Mutation of 5 amino acids on the dynein binding face of the yeast Lis1 β-propeller disrupts the interaction between the dynein motor domain and Lis1^7,28^. We made the equivalent mutations in human Lis1 (“Lis1-5A”; Fig. 2d). We first asked if Lis1-5A could enhance the velocity of activated dynein complexes. We focused on dynein/ dynactin complexes activated by BicD2-S for these experiments, as Lis1 had the greatest effect on these complexes (Fig. 1). We found that 300 nM Lis1-5A could still enhance the velocity of dynein/ dynactin/ BicD2-S complexes. We hypothesized that the mutations in Lis1-5A might not completely disrupt the dynein interaction. Thus, we lowered the concentration of Lis1 and Lis1-5A to 24 nM and found that under these conditions Lis1-5A increased velocity significantly less than wild type Lis1 (Fig. 2e). Next, we asked if the Lis1-5A mutant could increase the percentage of processive runs for dynein/ dynactin/ Hook3 complexes. We observed significantly fewer processive runs of dynein/ dynactin/ Hook3 complexes in the presence of Lis1-5A compared to wild type Lis1 (Fig. 2f) and the velocity of these complexes was no longer increased (Supplementary Fig. 2a). A high-resolution structure of the human dynein/ Lis1 complex will be required to map the Lis1/ dynein binding sites in detail.

Since Lis1 is a dimer, we wondered whether Lis1’s role in forming activated complexes required dimerization. To test this, we purified monomeric human Lis1, which lacks its amino-terminal dimerization domain. Our data indicate that dimerization is not required for Lis1 to increase dynein velocity: we found that monomeric Lis1 could still increase complex formation at the same molar ratio of dynein to Lis1 β-propellers (Fig. 2g).

As activated dynein complexes containing two dynein dimers are faster than those containing a single one^5^, we hypothesized that Lis1 may maximally activate dynein by promoting the recruitment of a second dynein dimer to the complex. First, we sought to determine if Lis1 enhances the formation of dynein/ dynactin complexes in vitro. We measured the formation of activated dynein complexes by mixing dynein and dynactin with a molar excess of each activating adaptor conjugated to magnetic beads via their carboxy-terminal HaloTags. We then quantified the percentage of dynein depleted from the supernatant after the beads were pelleted with a magnet (Fig. 3a). The presence of Lis1 increased the percentage of dynein depleted by each activating adaptor (Fig. 3b) and dynactin was required for this effect (Supplementary Fig. 3a,b).

**Figure 3.**
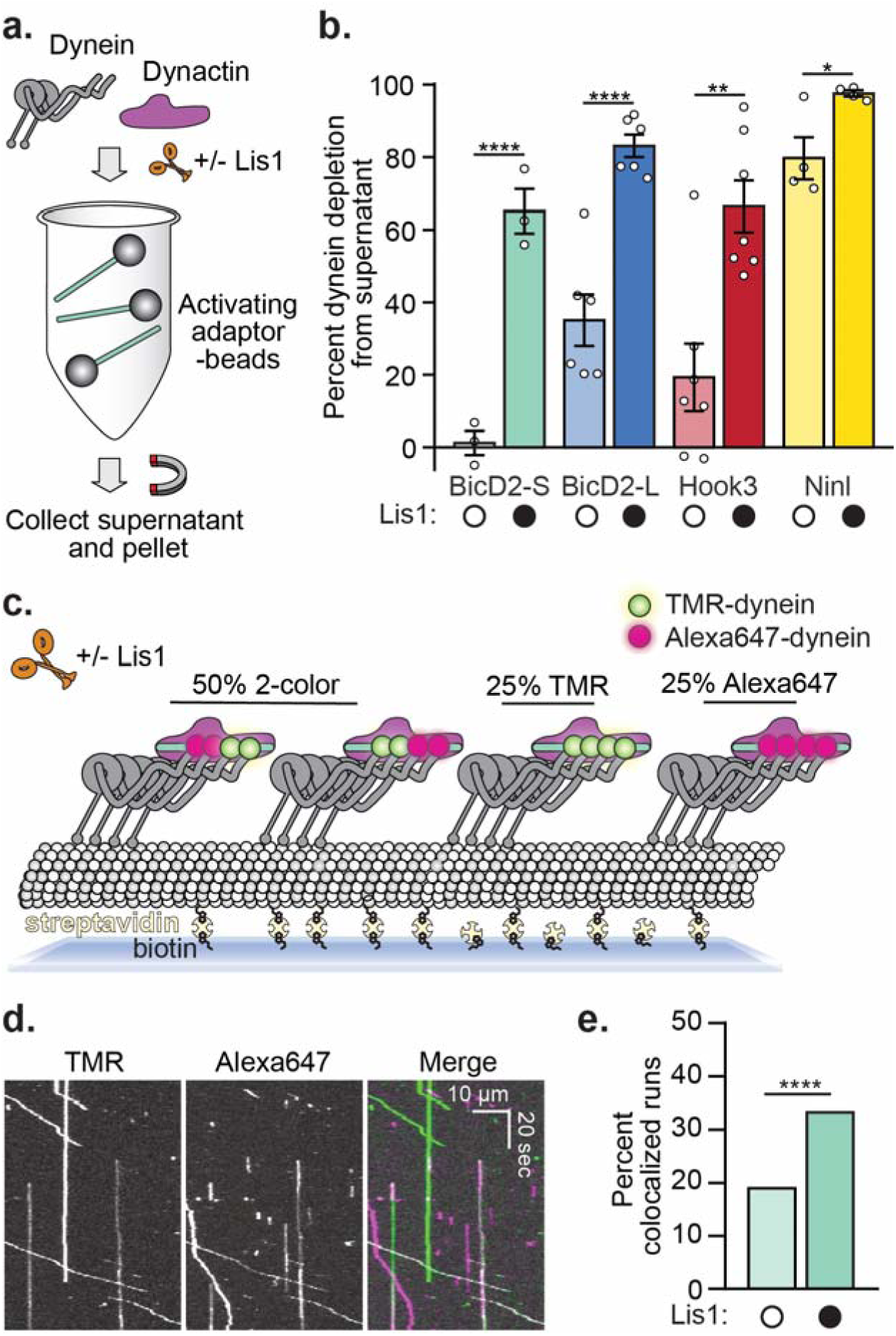
Lis1 recruits a second dynein dimer to dynein/ dynactin/ activating adaptor complexes. **a**. Schematic of the dynein depletion experiment performed in b. **b**. Percent dynein depletion (mean ± s.e.m.) by an activating adaptor conjugated to beads in the presence of dynactin and in the absence (white circles) or presence (black circles) of 150 nM Lis1. The activating adaptors conjugated to beads are indicated. Statistical analysis was performed using a two-tailed unpaired t test; ****, p<0.0001; **, p=0.0008; *, p=0.0017; n (replicates pre condition) = 3 (BicD2-S), 6 (BicD2-L), 7 (Hook3), 4 (Ninl). **c**. Schematic depicting the maximum probability of forming various dynein/ dynactin/ BicD2-S complexes containing two dynein dimers. The maximum probability of colocalization was 47% due to our labeling efficiency. **d**. Representative kymographs showing the colocalization of TMR- and Alexa647-labeled dynein in moving activated dynein/ dynactin/ BicD2-S complexes. Each channel is shown separately (left and middle panels) and the merged TMR- and Alexa647-channels in pseudocolor (right panel). **e**. Percent two-color dynein/ dynactin/ activating adaptor runs in the absence (white circles) or presence (black circles) of 300 nM Lis1. Statistical analysis was performed using a chi-squared test; ****, p<0.0001; n = 342 (no Lis1), 516 (with Lis1).

To directly test if Lis1 promotes the recruitment of a second dynein dimer we performed a two-color single-molecule assay where we mixed equimolar amounts of dynein labeled (on the heavy chain subunit) with either TMR or Alexa647. In this assay, if all moving activated dynein complexes contain two dynein dimers, 50% of motile events will show co-localized dynein-TMR and dynein-Alexa647 (Fig. 3c). We found that the presence of Lis1 significantly increased the number of moving two-color activated dynein/ dynactin/ BicD2-S complexes (Fig. 3d,e). From these experiments, we conclude that Lis1 promotes the recruitment of a second dynein dimer to form maximally activated dynein complexes.

We wondered if Lis1 must remain bound to moving activated dynein complexes to sustain fast velocity. To address this, we sought to determine if TMR-labeled Lis1 comigrated with moving dynein/ dynactin/ BicD2-S complexes tagged with Alexa647 (on the dynein heavy chain subunit). For most of our earlier experiments we used 300 nM Lis1, a concentration that is too high to visualize single Lis1 molecules. Therefore, we lowered the Lis1 concentration to 50 nM, a concentration where Lis1 still increased the velocity of dynein/ dynactin/ BicD2-S complexes (Fig. 4a). When we performed single-molecule motility assays with TMR-Lis1 and Alexa647-dynein/ dynactin/ BicD2-S complexes we found that only 16.8 ± 1.9% of dynein runs comigrated with Lis1 and those runs that did comigrate moved significantly slower than those with no detectable Lis1 (Fig. 4b,c). On a few occasions we were able to capture the disappearance of the TMR-Lis1 signal coincided with an increase in speed of activated dynein/ dynactin/ BicD2-S runs (Fig. 4d). When we performed the same experiment with the Lis1-5A mutant, we found that it increased velocity significantly less than wild type Lis1 (Supplementary Fig. 4a). However, we observed no colocalization of Lis1-5A with dynein (Supplementary Fig. 4b). These results suggest that the presence of Lis1 is not required for sustained maximal velocity of activated dynein complexes.

**Figure 4.**
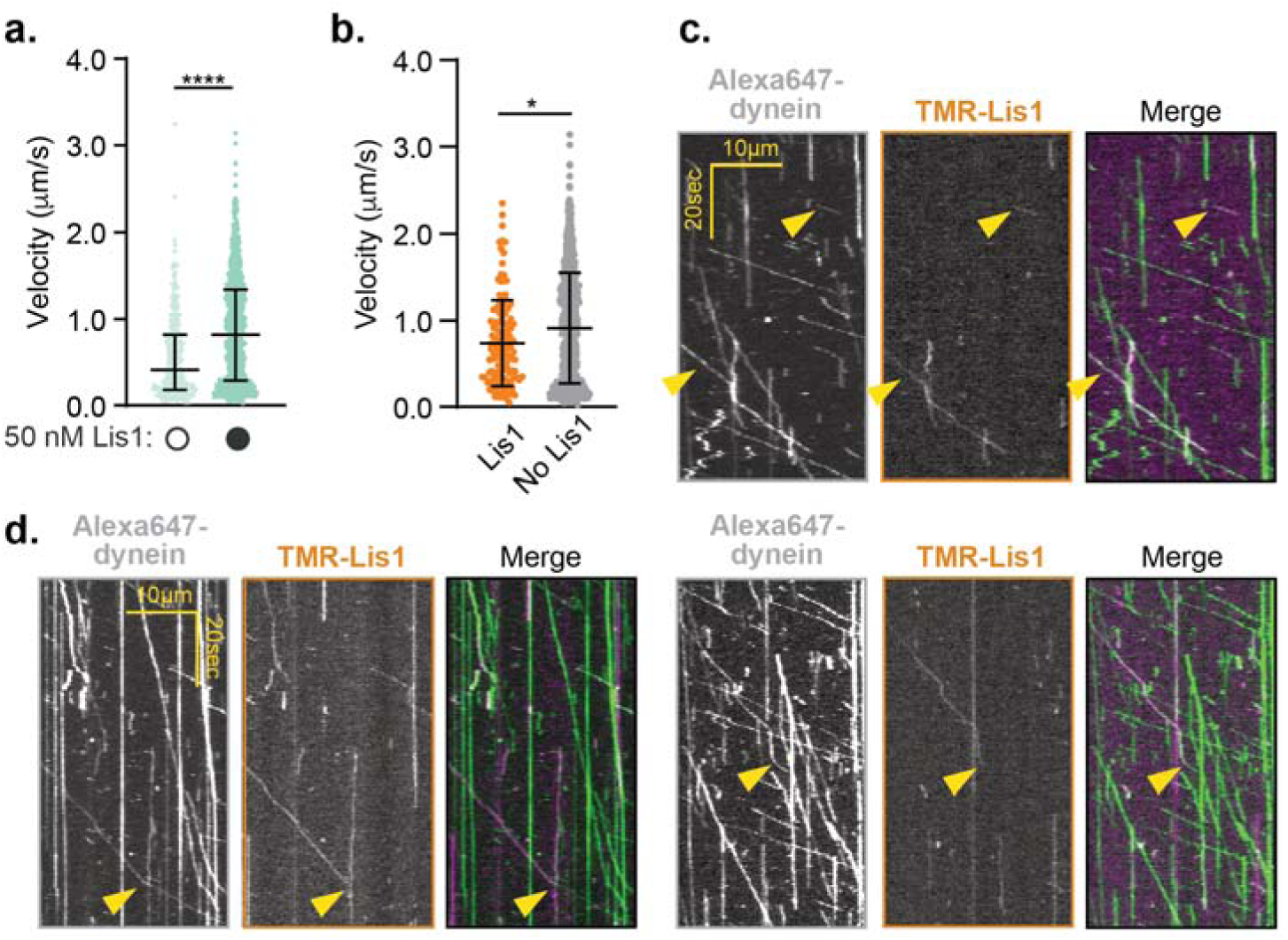
Lis1 is not required to sustain the maximal velocity of activated dynein complexes. **a**. Velocity of dynein/ dynactin/ BicD2-S complexes in the absence (white circles) or presence (black circles) of 50 nM TMR-Lis1. The median and interquartile range are shown. Statistical analysis was performed using a Kruskal-Wallis test with Dunn’s multiple comparisons test; ****, p<0.0001; n (individual single molecule events) = 331 (no Lis1), 924 (with Lis1). **b**. Velocity of dynein-Alexa647/ dynactin/ BicD2-S complexes in the presence of 50 nM TMR-Lis1, either colocalized with TMR-Lis1 (orange) or without detectable TMR-Lis1 (grey). The median and interquartile range are shown. 16.8 ± 1.9% of the dynein runs colocalized with Lis1. Statistical analysis was performed using a two-tailed Mann-Whitney test; *, p=0.0011; n (individual single molecule events) = 136 (co-localized Lis1), 781 (no detectable Lis1). **c**. Representative kymographs of the dynein and Lis1 channels (left and middle panels) and the merged images in pseudocolor (right panel). **d**. Kymographs showing examples of dynein’s velocity changing upon loss of the Lis1 signal. The dynein and Lis1 channels are shown (left and middle panels) and the merged images in pseudocolor (right panel).

Together our data indicate that Lis1 plays a role in the formation of maximally activated dynein complexes. We next explored at which step(s) of the dynein assembly pathway Lis1 was acting (Fig. 1a). Specifically, because the site_ring_ Lis1 binding site on dynein is not accessible in Phi dynein (Fig. 5a), we wondered if Lis1 could shift the equilibrium of Phi to Open dynein. In agreement with this idea, we found that Lis1 had a higher affinity for a dynein mutant that does not form the Phi conformation (K1610E and R1567E^12^, “Open dynein”; Fig. 5b). Next, we asked if Lis1 influenced dynein/ dynactin/ activating adaptor complex formation or velocity when the dynein used was the Open dynein mutant. We first examined Open dynein complexes in the absence of Lis1 and found that these dynein/ dynactin/ BicD2-S complexes were more likely to form and moved faster compared to complexes containing wild type dynein (Fig. 5c,d). We then asked if Lis1 affected activated complexes containing Open dynein. We found that Lis1 further increased complex formation and velocity of open dynein/dynactin/BicD2-S. Lis1 also increased the percentage of two dynein complexes when Open dynein was used (Fig. 5e).

**Figure 5.**
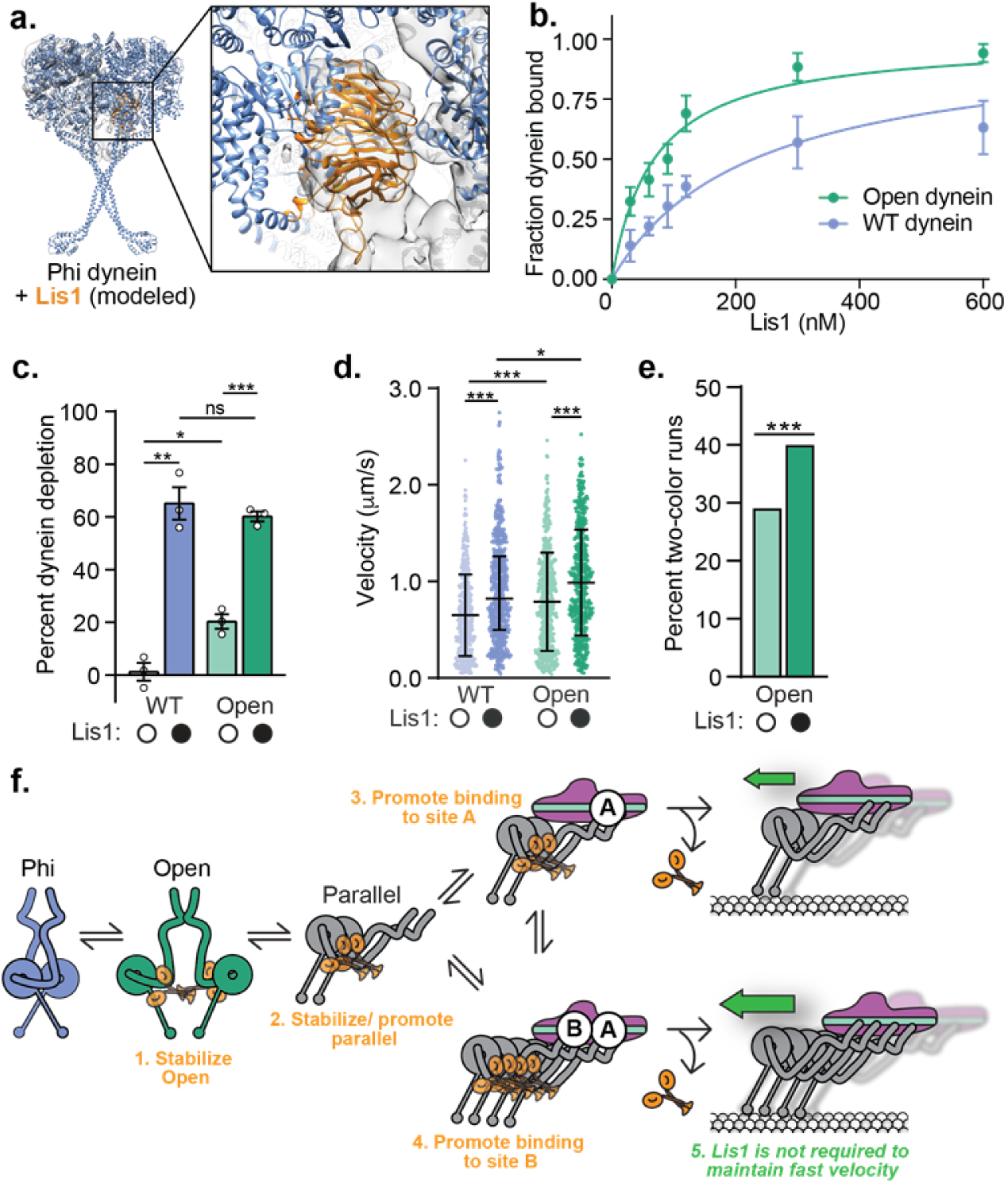
Lis1 is required to maximally activate Open dynein/ dynactin/ BicD2-S complexes. **a**. One of the dynein protomers in the Phi conformation (PDB: 5NVU) was aligned to the structure of yeast dynein (AAA3-Walker B) bound to Lis1 in the presence of ATP-vanadate (PDB: 5VLJ). The inset, which shows the cryo-EM map for the yeast structure with Lis1 docked at its AAA3/4 binding site (site_ring_), highlights the steric incompatibility between the Phi conformation and binding of Lis1 at AAA3/4. **b**. Determination of the binding affinity of Lis1 for wild type dynein (blue, K_d_ = 144 nM ± 25) and Open dynein (green, K_d_ = 80 nM ± 8.1). **c**. Percent (mean ± s.e.m.) depletion of WT dynein (blue) and Open dynein (green) by BicD2-S conjugated to beads in the absence (white circles) or presence (black circles) of 150 nM Lis1. Data with WT dynein in the presence and absence of Lis1 is also presented in Fig. 3b. Statistical analysis was performed using a two-tailed unpaired t test with Bonferroni corrected significance levels for two comparisons; ***, p=0.0003; **, p=0.0008; *, p=0.0118; ns, p=0.4857; n = 3 replicates per condition. **d**. Velocity of dynein/ dynactin/ BicD2-S complexes with wild type dynein (blue) and Open dynein (green) in the absence (white circles) or presence (black circles) of 300 nM Lis1. The median and interquartile range are shown. Data with WT dynein with and without Lis1 was also presented in Fig. 1f. Statistical analysis was performed using a two-tailed unpaired t test with Bonferroni corrected significance levels for two comparisons; ***, p<0.0001; *, p=0.0241; n (individual single molecule events) = 506 (WT dynein no Lis1), 569 (WT dynein with Lis1), 517 (Open dynein no Lis1), 507 (Open dynein with Lis1). **e**. Percent two-color colocalized runs (mean ± s.e.m.) with activated dynein complexes with Open dynein in the absence (white circles) or presence (black circles) of 300 nM Lis1. Statistical analysis was performed using a chi-squared test; ***, p=0.0010; n = 395 (no Lis1), 435 (with Lis1). **f**. Model for the roles of Lis1 in forming activated dynein complexes. See text for details.

Together our work suggests that Lis1 promotes the formation of activated dynein complexes reconstituted with purified human components. Experiments in human cells^18^, *Drosophila* embryos^24^, and *Xenopus* extracts^41^ have also shown that Lis1 is required for the interaction of dynein and dynactin with either its cargos or dynactin. Based on our data, our current model of dynein activation by Lis1 consists of three elements (Fig. 5f).

First, Lis1 may promote the Open dynein conformation. We hypothesize that this could occur by Lis1 shifting the equilibrium between Phi and Open dynein by binding to dynein at site_ring_ (Fig. 5f, step 1). We propose this based on our data showing that human Lis1 binds dynein at site_ring_ (which is inaccessible in Phi dynein, Fig. 5a), and our data showing that Lis1 has a higher affinity for dynein that cannot form the Phi particle. An assay that can report on both the Phi and Open conformation will be required to directly test this.

Second, Lis1 promotes or stabilizes the Parallel dynein conformation. We hypothesize that Lis1 binding to Open dynein favors a dynein conformation that leads to the assembly of the fully activated dynein complex (Fig. 5f, steps 2-4). Our data showing that the Open dynein mutant is further activated by Lis1 supports this idea. While we cannot rule out that some percentage of Open dynein molecules still adopt a partially autoinhibited conformation (that could be relieved by Lis1 as proposed above), we think this is unlikely given previous EM studies showing that the Open dynein mutant does not form Phi particles^12^. Our data also provide mechanistic insight for how Lis1 promotes complex formation. Our cryo-EM data shows that human Lis1, like yeast Lis1^7^, interacts with dynein at two sites on its motor domain. A high-resolution structure of human dynein bound to Lis1 and mutagenesis of both sites on dynein will be required to determine the role of each binding site in complex formation. However, because the Lis1-5A mutant is defective in supporting complex formation, and in yeast Lis1-5A mutants have a similar phenotype to site_ring_ mutants^7,28^, we hypothesize that binding at this site will be important for activated complex assembly. Importantly, because we found that monomeric Lis1 still influenced activated complex formation, Lis1 is likely not functioning by cross-linking adjacent dynein dimers (dynein A and B).

Third, Lis1 is not required for sustained dynein velocity. We hypothesize that Lis1 facilitates the formation of dynein/ dynactin/ activator complexes, but dissociates from moving complexes (Fig. 5f, step 5). Complex formation may occur at microtubule plus ends as previous in vitro studies showed that Lis1 increases the binding of activated dynein complexes to microtubule plus ends^42^. This component of our model is based on our data showing that most moving dynein complexes do not remain bound to Lis1 and those that do, move slower. This is in agreement with studies in *Aspergillus nidulans* showing that Lis1 is required for the initiation of endosome motility, but does not colocalize with moving endosomes^43^. We propose that filamentous fungi like *A. nidulans*, which have Hook-related activating adaptors that are required for dynein-based motility of endosomes^44,45^, also use Lis1 to promote activated dynein complex formation. Our model may also be conserved in yeast. While *S. cerevisiae* dynein is processive in vitro in the absence of dynactin or an activating adaptor^10^, Lis1 is required for dynein/ dynactin localization to its site of activation (the cortex) in vivo^21,46^. In yeast both dynein^47^ and Num1^48^, a candidate activating adaptor^49,50^ localize to the cortex, while a stable pool of Lis1 is not observed there^21,51^.

Finally, we have shown that proteins representing three distinct families of activating adaptors, BicD2, Hook3, and Ninl, all require Lis1 for full activation. This raises the possibility that many activated dynein complexes in cells may consist of two dynein dimers, dynactin, and an activating adaptor. We hypothesize that related activating adaptors or candidate activating adaptors will also require Lis1 to maximally activate dynein/ dynactin. In humans there are three additional BicD family members (BicD1, BicDL1, and BicDL2), two additional Hook family members (Hook1 and Hook2), and additional proteins that share a similar domain structure in their dynein/ dynactin binding regions to the Hook proteins (CCDC88A, CCDC88B, and CCDC88C) or Ninl (Nin, Rab11-FIP3, CRACR2a, Rab 44, and Rab 45)^2,9,52^. Thus, we predict that Lis1 will play a role in the cell biological processes that all of these activating adaptors or candidate activating adaptors facilitate.

## Acknowledgements

We thank Jenna Christensen and John Salogiannis for comments on the manuscript, the Nikon Imaging Center at UC San Diego where we collected data and received help with image analysis, and the UC San Diego cryo-EM facility, where cryo-EM data was collected. We also thank the Physics Computing Facility for IT support. SRP is supported by HHMI and NIH R01GM121772. Funding from the HHMI/Simons Faculty Scholars Program and R01GM107214 funded parts of this work. AEL is supported by R01GM107214, ZMH by an NSF graduate research fellowship DGE1144152, JPG by Molecular Biophysics Training Grant, NIH Grant T32 GM008326, RWB is a Damon Runyon Fellow supported by the Damon Runyon Cancer Research Foundation DRG-#2285-17, and MED by NIH K99GM127757 and previously by a Jane Coffin Childs Memorial Fund Postdoctoral Fellowship 61-1552-T.

## Author Contributions

ZMH, JPG, MED, AEL and SRP designed the experiments. ZMH, JPG, MED and RWB performed the experiments. ZMH, PJG, MED, AEL and SRP wrote the manuscript. All authors interpreted the data and reviewed and edited the manuscript.

---

**Fig. s1.**
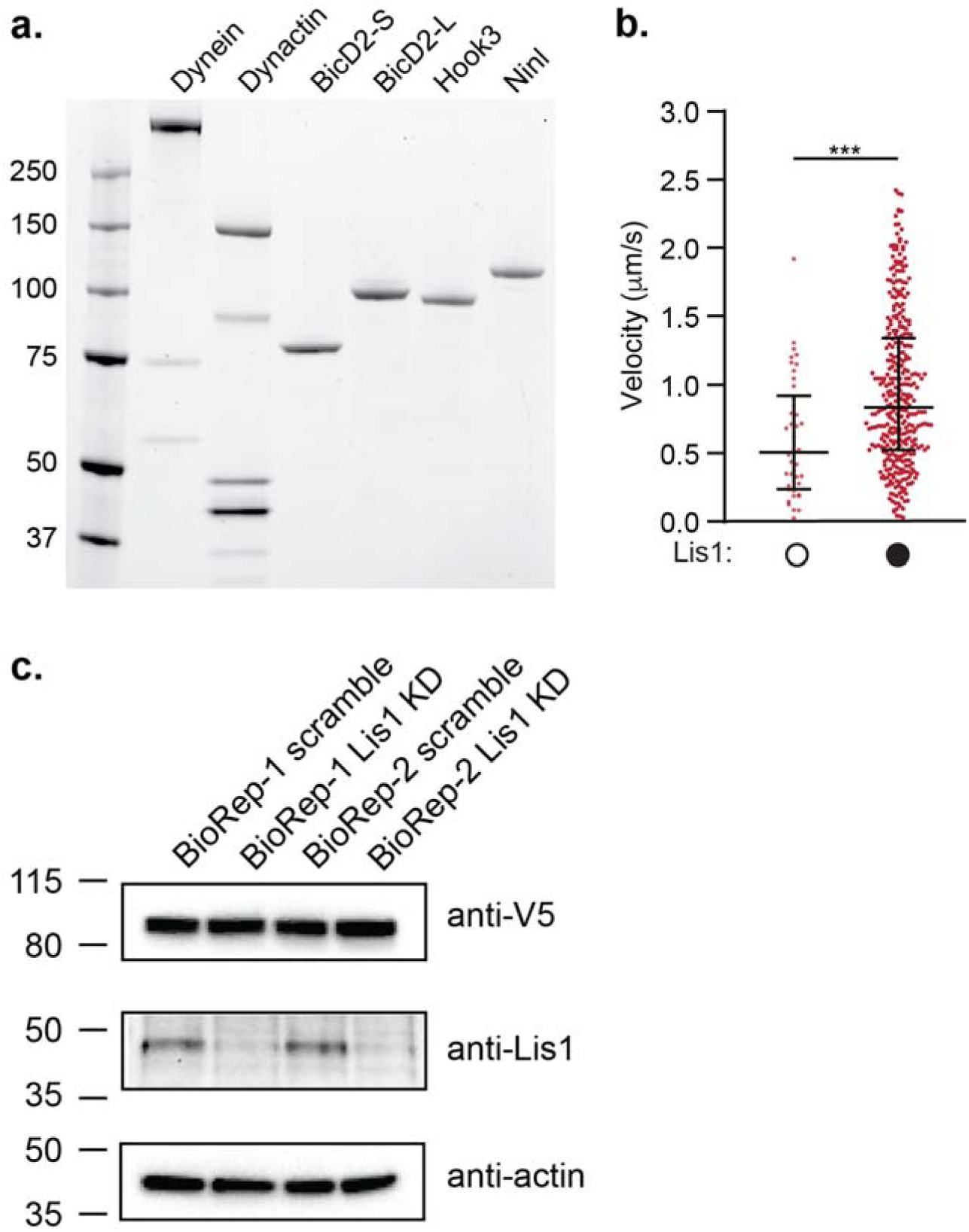
Supplemental figure relating to figure 1. **a**. SDS-PAGE gel stained with Sypro Red of human dynein, dynactin and the activating adaptors BicD2-S (aa 25-398), BicD2-L (aa 1-598), Hook3 (aa 1-552), and Ninl (aa 1-702) used here. The dynein heavy chain was tagged with the SNAP tag, the dynactin subunit p62 was tagged with the HaloTag, and each activating adaptor was tagged with the HaloTag. The dynein light chains are too small to be seen on this low percentage gel. **b**. Velocity of dynein/ dynactin/ Hook3 complexes in a higher salt buffer in the absence (white circles) or presence (black circles) of Lis1. The median and interquartile range is shown. Statistical analysis was performed using Kruskal-Wallis test with Dunn’s multiple comparisons test; ***, p=0.0004; n (individual single molecule events) = 42 (no Lis1), 353 (with Lis1). **c**. SDS-PAGE gel and immunoblots of cell lysates from human U2OS cells transfected with PEX3-mEMerald-FKBP and BicD2-S-V5-FRB constructs with scramble or Lis1 siRNA knockdown. An anti-V5 antibody detects BicD2-S-V5-FRB, an anti-Lis1 antibody assesses the efficiency of Lis1 knockdown, and an anti-actin antibody serves as a loading control. Two bio-replicates (biorep 1 and 2) are shown.

**Fig. s2.**
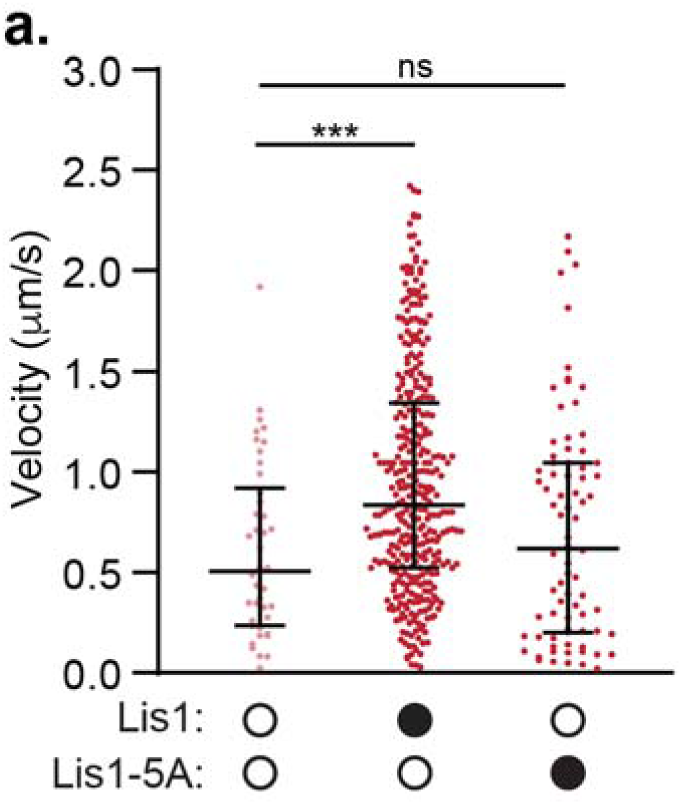
Supplemental figure relating to figure 2. **a**. Velocity of dynein/ dynactin/ Hook3 complexes in the presence of a higher salt buffer (60 mM KOAc versus 30 mM KOAc in our standard motility buffer) in the absence (white circles) or presence (black circles) of 300 nM Lis1 or Lis1-5A. The data in the presence and absence of WT Lis1 was also presented in Fig. S1b. The median and interquartile range are shown. Statistical analysis was performed using Kruskal-Wallis test with Dunn’s multiple comparisons test; ***, p=0.0004; ns, p>0.9999; n (individual single molecule events) = 42 (no Lis1), 353 (with Lis1), 78 (with Lis1-5A).

**Fig. s3.**
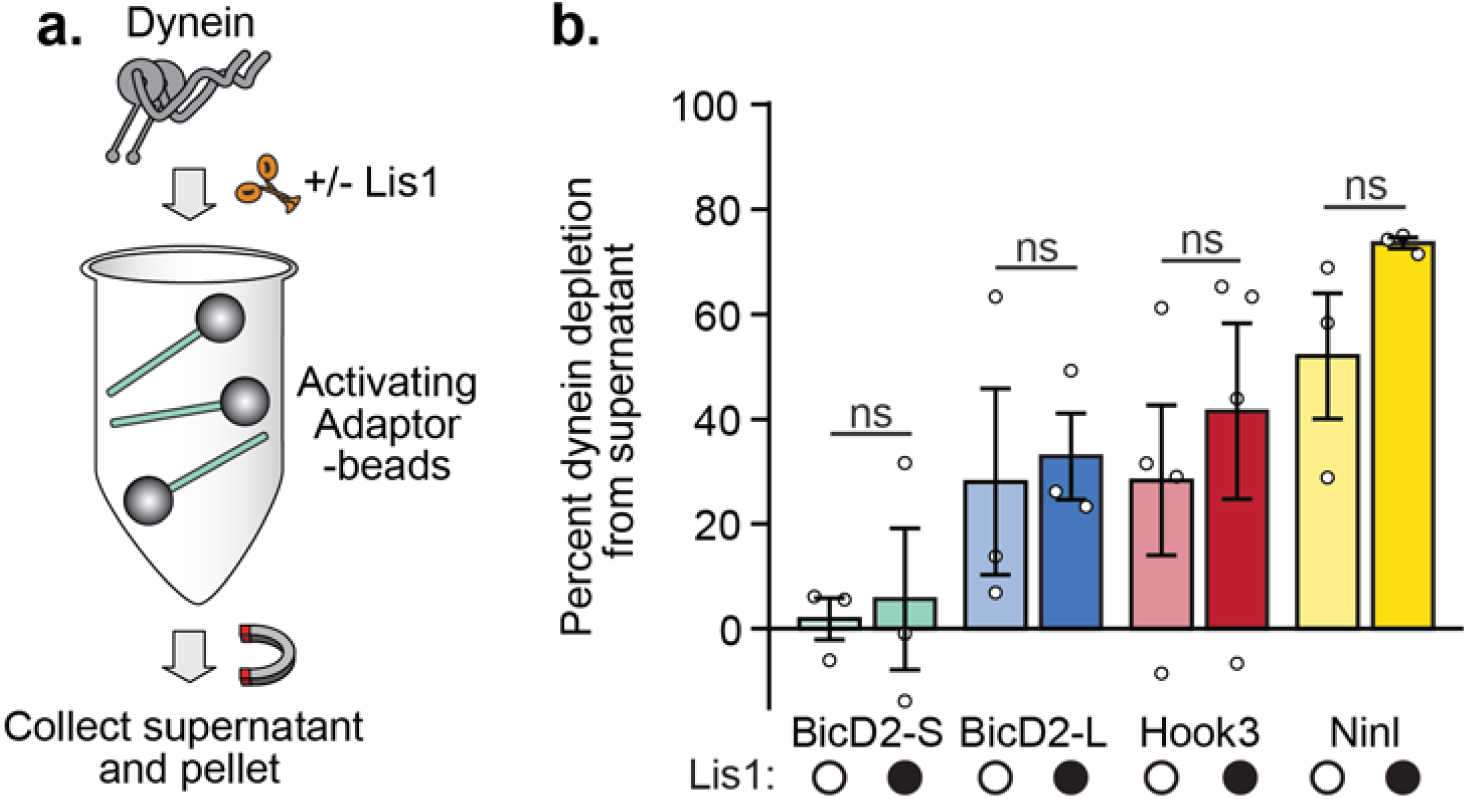
Supplemental figure relating to figure 3. **a**. Schematic of the dynein depletion experiment performed in b. **b**. Percent dynein depletion (mean ± s.e.m.) by an activating adaptor conjugated to beads in the absence of dynactin and in the absence (white circles) or presence (black circles) of 150 nM Lis1. The activating adaptors conjugated to beads are indicated. Statistical analysis was performed using a two-tailed unpaired t test; all comparisons were not significant (ns): p=0.8026 (BicD2-S), p=0.8152 (BicD2-L), p=0.5715 (Hook3); p=0.1472 (Ninl); n = 3 (BicD2-S, BicD2-L, Ninl) or 4 (Hook3) replicates.

**Fig. s4.**
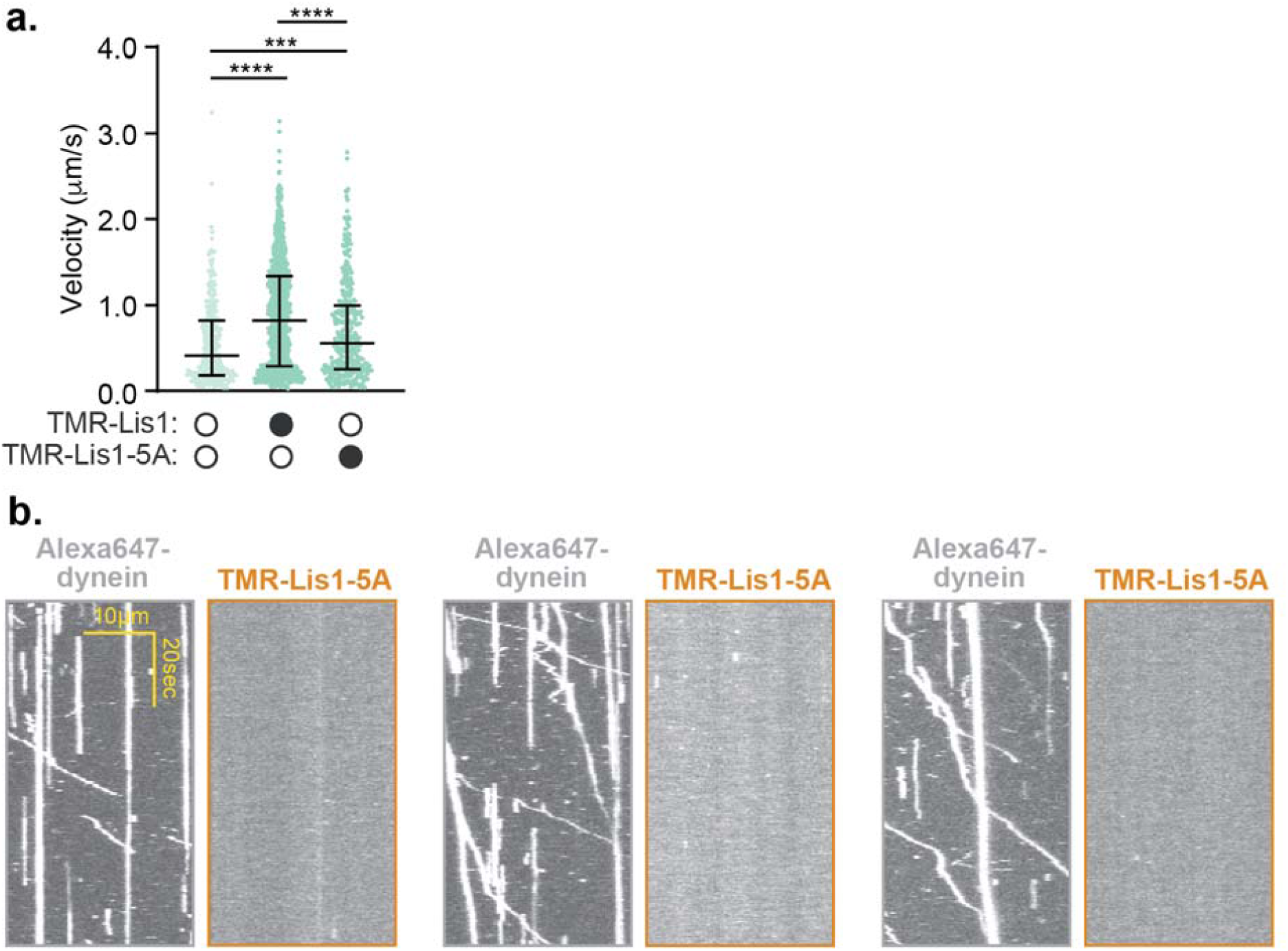
Supplemental figure relating to figure 4. **a**. Velocity of dynein/ dynactin/ BicD2-S complexes in the absence (white circles) or presence (black circles) of 50 nM TMR-Lis1 or TMR-Lis1-5A. The median and interquartile range are shown. Statistical analysis was performed using a Kruskal-Wallis test with Dunn’s multiple comparisons test; ****, p<0.0001; ***, p=0.0009; n (individual single molecule events) = 331 (no Lis1), 924 (WT Lis1), 378 (Lis1-5A). **b**. Single-molecule motility assays with dynein-Alexa647/ dynactin/ BicD2-S complexes in the presence of 50 nM TMR-Lis1-5A. Three pairs of representative kymographs are shown with the dynein channel (left) and the Lis1 channel (right).

## Supplementary Video Legends

### Video s1

Human U2OS cells co-transfected with PEX3-mEMerald-FKBP and BicD2-S-V5-FRB constructs and a scrambled siRNA. mEmerald dynamics are visualized before addition of rapalog. Frames were taken every 500 msec for 2 min. Video frame rate is 20 frames/sec.

### Video s2

Human U2OS cells co-transfected with PEX3-mEMerald-FKBP and BicD2-S-V5-FRB constructs and a scrambled siRNA. mEmerald dynamics are visualized after addition of rapalog. Orange arrows indicate examples of motile peroxisomes. Frames were taken every 500 msec for 2 min. Video frame rate is 20 frames/sec.

### Video s3

Human U2OS cells co-transfected with PEX3-mEMerald-FKBP and BicD2-S-V5-FRB constructs and Lis1 siRNA. mEmerald dynamics are visualized before addition of rapalog. Frames were taken every 500 msec for 2 min. Video frame rate is 20 frames/sec.

### Video s4

Human U2OS cells co-transfected with PEX3-mEMerald-FKBP and BicD2-S-V5-FRB constructs and Lis1 siRNA. mEmerald dynamics are visualized after addition of rapalog. Orange arrows indicate examples of motile peroxisomes. Frames were taken every 500 msec for 2 min. Video frame rate is 20 frames/sec.

## METHODS

### Cloning, plasmid construction, and mutagenesis

The pDyn1 plasmid (the pACEBac1 expression vector containing insect cell codon optimized dynein heavy chain (*DYNC1H1*) fused to a His-ZZ-TEV tag on the amino-terminus and a carboxy-terminal SNAPf tag (New England Biolabs)) and the pDyn2 plasmid (the pIDC expression vector with codon optimized *DYNC1I2, DYNC1LI2, DYNLT1, DYNLL1*, and *DYNLRB1*) were recombined in vitro with a Cre recombinase (New England Biolabs) to generate the pDyn3 plasmid. The presence of all six dynein chains was verified by PCR. pDyn1, pDyn2 and the pFastBac plasmid with codon-optimized human full-length Lis1 (*PAFAH1B1*) fused to an amino-terminal His-ZZ-TEV tag and pFastBac containing human dynein monomer (amino acids 1320-4646 of DYNC1H1) were gifts from Andrew Carter (LMB-MRC, Cambridge, UK). BicD2 constructs were amplified from a human cDNA library generated from RPE1 cells and the other activating adaptor constructs were obtained as described previously^34^. Activating adaptors were fused to a ZZ-TEV-HaloTag (Promega) on the amino-terminus and inserted into a pET28a expression vector. All additional tags were added via Gibson assembly and all mutations and truncations were made via site-directed mutagenesis (Agilent). For rapalog induced motility in cells, HaloTag-BicD2-S was cloned into the pcDNA5 backbone with a carboxy-terminal V5 epitope tag fused to FRB. The peroxisome tag PEX3 was cloned into pcDNA5 with a carboxy-terminal mEmerald fluorescent protein and FKBP.

### Protein expression and purification

Human full-length dynein, human dynein monomer, and human Lis1 constructs were expressed in Sf9 cells as described previously^3,32^. Briefly, the pDyn3 plasmid containing the human dynein genes or the pFastBac plasmid containing full-length Lis1 or dynein monomer was transformed into DH10EmBacY chemically competent cells with heat shock at 42°C for 15 seconds followed by incubation at 37°C for 5 hours in S.O.C media (Thermofisher scientific). The cells were then plated on LB-agar plates containing kanamycin (50 μg/ml), gentamicin (7 μg/ml), tetracyclin (10 μg/ml), BluoGal (100 μg/ml) and IPTG (40 μg/ml) and positive clones were identified by a blue/white color screen after 48 hours. For full-length human dynein constructs, white colonies were additionally tested for the presence of all six dynein genes using PCR. These colonies were then grown overnight in LB medium containing kanamycin (50 μg/ml), gentamicin (7 μg/ml) and tetracyclin (10 μg/ml) at 37°C. Bacmid DNA was extracted from overnight cultures using an isopropanol precipitation method as described previously ^12^. 2mL of Sf9 cells at 0.5×10^6^ cells/mL were transfected with 2µg of fresh bacmid DNA and FuGene HD transfection reagent (Promega) at a 3:1 transfection reagent to DNA ratio according to the manufacturer’s instructions. After three days, the supernatant containing the “V0” virus was harvested by centrifugation at 200 x g for 5 minutes at 4°C. To generate “V1”, 1 mL of the V0 virus was used to transfect 50mL of Sf9 cells at 1×10^6^ cells/mL. After three days, the supernatant containing the V1 virus was harvested by centrifugation at 200 x g for 5 minutes at 4°C and stored in the dark at 4°C until use. For protein expression, 4 mL of the V1 virus were used to transfect 400 mL of Sf9 cells at 1×10^6^ cells/mL. After three days, the cells were harvested by centrifugation at ×3000 ×g for 10 minutes at 4°C. The pellet was resuspended in 10 mL of ice-cold PBS and pelleted again. The pellet was flash frozen in liquid nitrogen and stored at −80°C.

Protein purification steps were done at 4°C unless otherwise indicated. Full-length dynein and dynein monomer were purified from frozen Sf9 pellets transfected with the V1 virus as described previously^3^. Frozen cell pellets from a 400 mL culture were resuspended in 40 mL of Dynein-lysis buffer (50 mM HEPES [pH 7.4], 100 mM sodium chloride, 1 mM DTT, 0.1 mM Mg-ATP, 0.5 mM Pefabloc, 10% (v/v) glycerol) supplemented with 1 cOmplete EDTA-free protease inhibitor cocktail tablet (Roche) per 50 mL and lysed using a Dounce homogenizer (10 strokes with a loose plunger and 15 strokes with a tight plunger). The lysate was clarified by centrifuging at 183,960 x g for 88 min in Type 70 Ti rotor (Beckman). The clarified supernatant was incubated with 4 mL of IgG Sepharose 6 Fast Flow beads (GE Healthcare Life Sciences) for 3-4 hours on a roller. The beads were transferred to a gravity flow column, washed with 200 mL of Dynein-lysis buffer and 300 mL of TEV buffer (50 mM Tris–HCl [pH 8.0], 250 mM potassium acetate, 2 mM magnesium acetate, 1 mM EGTA, 1 mM DTT, 0.1 mM Mg-ATP, 10% (v/v) glycerol). For fluorescent labeling of carboxy-terminal SNAPf tag, dynein-coated beads were labeled with 5 µM SNAP-Cell-TMR (New England Biolabs) in the column for 10 min at room temperature and unbound dye was removed with a 300 mL wash with TEV buffer at 4°C. The beads were then resuspended and incubated in 15 mL of TEV buffer supplemented with 0.5 mM Pefabloc and 0.2 mg/mL TEV protease (purified in the Reck-Peterson lab) overnight on a roller. The supernatant containing cleaved proteins was concentrated using a 100K MWCO concentrator (EMD Millipore) to 500 µL and purified via size exclusion chromatography on a TSKgel G4000SWXL column (TOSOH Bioscience) with GF150 buffer (25 mM HEPES [pH7.4], 150 mM KCl, 1mM MgCl2, 5 mM DTT, 0.1 mM Mg-ATP) at 1 mL/min. The peak fractions were collected, buffer exchanged into a GF150 buffer supplemented with 10% glycerol, concentrated to 0.1-0.5 mg/mL using a 100K MWCO concentrator (EMD Millipore) and flash frozen in liquid nitrogen.

Lis1 constructs were purified from frozen cell pellets from 400 mL culture. Lysis and clarification steps were similar to full-length dynein purification except Lis1-lysis buffer (30 mM HEPES [pH 7.4], 50 mM potassium acetate, 2 mM magnesium acetate, 1 mM EGTA, 300 mM potassium chloride, 1 mM DTT, 0.5 mM Pefabloc, 10% (v/v) glycerol) supplemented with 1 cOmplete EDTA-free protease inhibitor cocktail tablet (Roche) per 50 mL was used. The clarified supernatant was incubated with 0.5 mL of IgG Sepharose 6 Fast Flow beads (GE Healthcare Life Sciences) for 2-3 hours on a roller. The beads were transferred to a gravity flow column, washed with 20 mL of Lis1-lysis buffer, 100 mL of modified TEV buffer (10 mM Tris–HCl [pH 8.0], 2 mM magnesium acetate, 150mM potassium acetate, 1 mM EGTA, 1 mM DTT, 10% (v/v) glycerol) supplemented with 100 mM potassium acetate, and 50 mL of modified TEV buffer. For fluorescent labeling of Lis1 constructs with amino-terminal HaloTags, Lis1-coated beads were labeled with 200 µM Halo-TMR (Promega) for 2.5 hours at 4°C on a roller and the unbound dye was removed with a 200 mL wash with modified TEV buffer supplemented with 250 mM potassium acetate. Lis1 was cleaved from IgG beads via incubation with 0.2 mg/mL TEV protease overnight on a roller. The cleaved Lis1 was filtered by centrifuging with an Ultrafree-MC VV filter (EMD Millipore) in a tabletop centrifuge and flash frozen in liquid nitrogen.

Dynactin was purified from stable HEK293-T cell lines expressing p62-Halo-3xFlag as described previously^34^. Briefly, frozen pellets collected from 160 × 15cm plates were resuspended in 80 mL of Dynactin-lysis buffer (30 mM HEPES [pH 7.4], 50 mM potassium acetate, 2 mM magnesium acetate, 1 mM EGTA, 1 mM DTT, 10% (v/v) glycerol) supplemented with 0.5 mM Mg-ATP, 0.2% Triton X-100 and 1 cOmplete EDTA-free protease inhibitor cocktail tablet (Roche) per 50 mL and rotated slowly for 15 min. The lysate was clarified by centrifuging at 66,000 x g for 30 min in Type 70 Ti rotor (Beckman). The clarified supernatant was incubated with 1.5 mL of anti-Flag M2 affinity gel (Sigma-Aldrich) overnight on a roller. The beads were transferred to a gravity flow column, washed with 50 mL of wash buffer (Dynactin-lysis buffer supplemented with 0.1 mM Mg-ATP, 0.5 mM Pefabloc and 0.02% Triton X-100), 100 mL of wash buffer supplemented with 250 mM potassium acetate, and again with 100 mL of wash buffer. For fluorescent labeling the HaloTag, dynactin-coated beads were labeled with 5 µM Halo-JF646 (Janelia) in the column for 10 min at room temperature and the unbound dye was washed with 100 mL of wash buffer at 4°C. Dynactin was eluted from beads with 1 mL of elution buffer (wash buffer with 2 mg/mL of 3xFlag peptide). The eluate was collected, filtered by centrifuging with Ultrafree-MC VV filter (EMD Millipore) in a tabletop centrifuge and diluted to 2 mL in Buffer A (50 mM Tris-HCl [pH 8.0], 2 mM MgOAc, 1 mM EGTA, and 1 mM DTT) and injected onto a MonoQ 5/50 GL column (GE Healthcare and Life Sciences) at 1 mL/min. The column was pre-washed with 10 CV of Buffer A, 10 CV of Buffer B (50 mM Tris-HCl [pH 8.0], 2 mM MgOAc, 1 mM EGTA, 1 mM DTT, 1 M KOAc) and again with 10 CV of Buffer A at 1 mL/min. To elute, a linear gradient was run over 26 CV from 35-100% Buffer B. Pure dynactin complex eluted from ∼75-80% Buffer B. Peak fractions containing pure dynactin complex were pooled, buffer exchanged into a GF150 buffer supplemented with 10% glycerol, concentrated to 0.02-0.1 mg/mL using a 100K MWCO concentrator (EMD Millipore) and flash frozen in liquid nitrogen.

Activating adaptors containing amino-terminal HaloTags were expressed in BL-21[DE3] cells (New England Biolabs) at OD 0.4-0.6 with 0.1 mM IPTG for 16 hr at 18°C. Frozen cell pellets from 2 L culture were resuspended in 60mL of activator-lysis buffer (30 mM HEPES [pH 7.4], 50 mM potassium acetate, 2 mM magnesium acetate, 1 mM EGTA, 1 mM DTT, 0.5 mM Pefabloc, 10% (v/v) glycerol) supplemented with 1 cOmplete EDTA-free protease inhibitor cocktail tablet (Roche) per 50 mL and 1 mg/mL lysozyme. The resuspension was incubated on ice for 30 min and lysed by sonication. The lysate was clarified by centrifuging at 66,000 x g for 30 min in Type 70 Ti rotor (Beckman). The clarified supernatant was incubated with 2 mL of IgG Sepharose 6 Fast Flow beads (GE Healthcare Life Sciences) for 2 hr on a roller. The beads were transferred to a gravity flow column, washed with 100 mL of activator-lysis buffer supplemented with 150 mM potassium acetate and 50mL of cleavage buffer (50 mM Tris–HCl [pH 8.0], 150 mM potassium acetate, 2 mM magnesium acetate, 1 mM EGTA, 1 mM DTT, 0.5 mM Pefabloc, 10% (v/v) glycerol). The beads were then resuspended and incubated in 15 mL of cleavage buffer supplemented with 0.2 mg/mL TEV protease overnight on a roller. The supernatant containing cleaved proteins were concentrated using a 50K MWCO concentrator (EMD Millipore) to 1 mL, filtered by centrifuging with Ultrafree-MC VV filter (EMD Millipore) in a tabletop centrifuge, diluted to 2 mL in Buffer A (30 mM HEPES [pH 7.4], 50 mM potassium acetate, 2 mM magnesium acetate, 1 mM EGTA, 10% (v/v) glycerol and 1 mM DTT) and injected onto a MonoQ 5/50 GL column (GE Healthcare and Life Sciences) at 1 mL/min. The column was pre-washed with 10 CV of Buffer A, 10 CV of Buffer B (30 mM HEPES [pH 7.4], 1 M potassium acetate, 2 mM magnesium acetate, 1 mM EGTA, 10% (v/v) glycerol and 1 mM DTT) and again with 10 CV of Buffer A at 1 mL/min. To elute, a linear gradient was run over 26 CV from 0-100% Buffer B. The peak fractions containing Halo-tagged activating adaptors were collected and concentrated to using a 50K MWCO concentrator (EMD Millipore) to 0.2 mL. For fluorescent labeling the HaloTag, the concentrated peak fractions were incubated with 5 □M Halo-Alexa488 (Promega) for 10 min at room temperature. Unbound dye was removed by PD-10 desalting column (GE Healthcare and Life Sciences) according to the manufacturer’s instructions. The labeled activating adaptor sample was concentrated using a 50K MWCO concentrator (EMD Millipore) to 0.2 mL, diluted to 0.5 mL in GF150 buffer and further purified via size exclusion chromatography on a Superose 6 Increase 10/300 GL column (GE Healthcare and Life Sciences) with GF150 buffer at 0.5 mL/min. The peak fractions were collected, buffer exchanged into a GF150 buffer supplemented with 10% glycerol, concentrated to 0.2-1 mg/mL using a 50K MWCO concentrator (EMD Millipore) and flash frozen in liquid nitrogen.

### Single-molecule TIRF microscopy

Single-molecule imaging was performed with an inverted microscope (Nikon, Ti-E Eclipse) equipped with a 100x 1.49 N.A. oil immersion objective (Nikon, Plano Apo) and a ProScan linear motor stage controller (Prior). The microscope was equipped with a LU-NV laser launch (Nikon), with 405 nm, 488 nm, 532 nm, 561 nm and 640 nm laser lines. The excitation and emission paths were filtered using appropriate single bandpass filter cubes (Chroma). For two-color colocalization imaging, the emission signals were further filtered and split using W-view Gemini image splitting optics (Hamamatsu). The emitted signals were detected with an electron multiplying CCD camera (Andor Technology, iXon Ultra 897). Illumination and image acquisition was controlled by NIS Elements Advanced Research software (Nikon).

Single-molecule motility and microtubule binding assays were performed in flow chambers assembled as described previously^53^ using the TIRF microscopy set up described above. Either biotin-PEG-functionalized coverslips (Microsurfaces) or No. 1-1/2 coverslips (Corning) sonicated in 100% ethanol for 10 min were used for the flow-chamber assembly. Taxol-stabilized microtubules with ∼10% biotin-tubulin and ∼10% fluorescent-tubulin (Alexa405-, 488-or 647-labeled) were prepared as described previously^29^. Flow chambers were assembled with taxol-stabilized microtubules by incubating sequentially with the following solutions, interspersed with two washes with assay buffer (30 mM HEPES [pH 7.4], 2 mM magnesium acetate, 1 mM EGTA, 10% glycerol, 1 mM DTT) supplemented with 20 µM Taxol in between: (1) 1 mg/mL biotin-BSA in assay buffer (3 min incubation); (2) 0.5 mg/mL streptavidin in assay buffer (3 min incubation) and (3) a fresh dilution of taxol-stabilized microtubules in assay buffer (3 min incubation). After flowing in microtubules, the flow chamber was washed twice with assay buffer supplemented with 1 mg/mL casein and 20 µM Taxol.

To assemble dynein-dynactin-activating adaptor complexes, purified dynein (10-20 nM concentration), dynactin and the activating adaptor were mixed at 1:2:10 molar ratio and incubated on ice for 10 min. These dynein-dynactin-activating adaptor complexes were then incubated with Lis1 or modified TEV buffer (to buffer match for experiments without Lis1) for 10 min on ice. Dynein alone was used instead of dynein-dynactin-activating adaptor complexes for the experiments with dynein alone. The mixtures of dynein, dynactin, activating adaptor and Lis1 were then flowed into the flow chamber assembled with taxol-stabilized microtubules. The final imaging buffer contained the assay buffer supplemented with 20 µM Taxol, 1 mg/mL casein, 71.5 mM β-mercaptoethanol, an oxygen scavenger system, and 1 mM Mg-ATP. The final concentration of dynein in the flow chamber was 0.5-1 pM for experiments with dynein-dynactin-activating adaptor complexes and 0.3-0.5 pM for dynein alone experiments. The final concentration of Lis1 was between 24 nM - 300 nM (as indicated in the main text) for experiments with unlabeled Lis1, and 50 nM for experiments with TMR-labeled Lis1. For single-molecule motility assays, microtubules were imaged first by taking a single-frame snapshot. Dynein and/or the activating adaptor labeled with fluorophores (TMR, Alexa647 or Alexa488) was imaged every 300 msec for 3 min. At the end, microtubules were imaged again by taking a snapshot to assess stage drift. Movies showing significant drift were not analyzed. Each sample was imaged no longer than 15 min. For single-molecule microtubule binding assays, the final imaging mixture containing dynein was incubated for an additional 5 min in the flow chamber at room temperature before imaging. After 5 min incubation, microtubules were imaged first by taking a single-frame snapshot. Dynein and/or activating adaptors labeled with fluorophores (TMR, Alexa647 or Alexa488) were imaged by taking a single-frame snapshot. Each sample was imaged at 4 different fields of view and there were between 5 and 10 microtubules in each field of view. In order to compare the effect of Lis1 on microtubule binding, the samples with and without Lis1 were imaged in two separate flow chambers made on the same coverslip on the same day with the same stock of polymerized tubulin as described previously^7^.

### Single-molecule motility assay analysis

Kymographs were generated from motility movies and dynein velocity was calculated from kymographs using ImageJ macros as described^54^. Only runs that were longer than 4 frames (1.2 s) were included in the analysis. Bright aggregates, which were less than 5% of the population, were excluded from the analysis. For two-color colocalization analysis, kymographs from each channel were generated and merged in ImageJ and the number of colocalized runs was determined manually. Data plotting and statistical analyses were performed in Prism8 (GraphPad).

### Single-molecule microtubule binding assay analysis

Intensity profiles of dynein or activating adaptor spots from a single-frame snapshot were generated over a 5-pixel wide line drawn perpendicular to the long axis of microtubules in ImageJ. Intensity peaks at least 2-fold higher than the neighboring background intensity were counted as dynein or activating adaptor spots bound to microtubules. Bright aggregates that were 5-fold brighter than the neighboring intensity peaks were not counted. The average binding density was calculated as the total number of dynein or activating adaptor spots divided by the total microtubule length in each snapshot. Normalized binding density was calculated by dividing by the average binding density of dynein or activating adaptor without Lis1 collected on the same coverslip (see above). Data plotting and statistical analyses were performed in Prism7 (GraphPad).

### Protein binding assays

To assess dynein/ dynactin complex formation, activating adaptors were first coupled to 15 µL of Magne HaloTag Beads (Promega) in 2 mL Protein Lo Bind Tubes (Eppendorf) using the following protocol. Beads were washed twice with 1 mL of GF150 without ATP supplemented with 10% glycerol and 0.1%NP40. Activating adaptors were diluted in this buffer to 75 nM. 25 uL of each diluted activating adaptor was added to the beads and gently shaken for one hour. 20 µL of supernatant were then analyzed via SDS-PAGE to confirm complete depletion of the activating adaptors. The activating adaptor-conjugated beads were washed once with 1 mL GF150 with 10% glycerol and 0.1% NP40 and once with 1mL of binding buffer (30 mM HEPES [pH 7.4], 2 mM magnesium acetate, 1 mM EGTA, 10% glycerol, 1 mM DTT, 1 mg/mL casein, 0.1% NP40, 1mM ADP) supplemented with 15.7 mM KCl and 8.3 mM KOAc. 10 nM dynein, 10nM dynactin and 150 nM Lis1 were diluted in binding buffer, which resulted in 15.7 mM KCl and 8.3 mM KOAc. For experiments lacking dynactin or Lis1 the protein dilutions were supplemented with equivalent amounts of their purification buffers. 25 µL of the dynein, dynactin and Lis1 mixture were added to the beads pre-bound with activating adaptors and gently agitated for 45 minutes. After incubation 20 µL of the supernatant was removed, and 6.67 µL of NuPAGE® LDS Sample Buffer (4X) and 1.33 µL of Beta-mercaptoethanol was added to each. The samples were boiled for 5 minutes before running on a 4-12% NuPAGE Bis-Tris gel at 4C. Depletion was determined using densitometry in ImageJ.

Lis1 binding curves were determined as above with minor variations. 25 µL of Magnet HaloTag Beads were used, and washed twice with 1 mL modified TEV buffer. 0, 30, 60, 90, 120, 300 and 600 nM Lis1 was bound to beads for one hour at ambient temperature. Beads were then washed with 1 mL of modified TEV buffer and 1 mL of binding buffer supplemented with 30 mM KCl and 6 mM KOAc. 10 nM of dynein was diluted in binding buffer supplemented with salt to 30 mM KCl and 6 mM KOAc. Binding and determination of depletion were carried out as above. Binding curves were fit in Prism7 (Graphpad) with a nonlinear regression for one site binding with Bmax set to 1.

### Cryo-EM sample preparation

A final concentration of 3.5 μM dynein monomer and 3.5 μM HaloTag-Lis1 were incubated in assay buffer supplemented with DTT, NP40, and ATPVO_4_ for 10-20 minutes before grids were prepared. Proteins were diluted and mixed such that the final salt and additive concentrations were 52.5 mM KCl, 20 mM KOAc, 4.8% glycerol, 5 mM DTT, 0.005% NP40, and 2.5 mM ATPVO_4_. 4 ul of sample was applied to UltraAuFoil R 1.2/1.3 300 Mesh grids (Electron Microscopy Sciences) that were glow discharged with 20 mA negative current for 30 sec. Grids were plunge-frozen in a Vitrobot Mark IV robot (FEI Company), maintained at 100% humidity and 4 °C.

### Cryo-EM data collection and image analysis

Data was collected on a Talos Arctica transmission electron microscope (FEI Company) operating at 200 keV with a K2 Summit direct electron detector (Gatan Inc). Dose-fractionated movies were collected in counting mode, with a final calibrated pixel size of 1.16 Å/pixel, a dose rate of ∼6 e^-^/pixel/sec, and a total dose of ∼60 e^-^/Å^2^. Leginon ^55^ was used for automated data collection and movies were processed on-the-fly using Appion^56^. Movie alignment was performed with MotionCor2^57^ defocus estimations were performed with CTFFIND4^58^, and particles were picked using DoG Picker^59^ 403,439 particles were extracted from 2,422 aligned, dose weighted micrographs in Relion-3^60^ with a box size of 288 × 288 pixels and binned by 2 for a final pixel size of 2.32 Å/pixel. The extracted particles were imported into cryoSPARC 2.4.2 for all subsequent analysis^61^. To generate the 2D-class averages shown in figure 2, two rounds of 2D classification were performed. In the first round, 2D classes containing clear density corresponding to the dynein ATPase ring and Lis1 (comprising 71,436 particles) were selected. In the second round, three classes, containing 22,621 total particles were selected for presentation in figure 2 (7,712 particles in the class on the left, 8,943 particles in the class in the middle, 6,506 particles in the class on the right).

To generate a model of human dynein bound to Lis1, we aligned human dynein-2 bound to ATP-vanadate (PDB: 4RH7^40^) with yeast dynein (AAA3-Walker B) in ATP-vanadate and Lis1 (PDB: 5VLJ^7^) using the part of the sequence that encompasses the two Lis1 binding sites in yeast dynein, from AAA3 until after the binding site for the second Lis1 in the stalk. We then deleted the dynein chain of 5VLJ and combined the remaining two copies of Lis1 with 4RH7. To highlight the densities corresponding to Lis1, we also generated 2D projections of human dynein-2 alone (PDB: 4RH7) in the same orientations as our experimental data.

### Peroxisome recruitment assay

Human U2OS cells were cultured in Dulbecco’s modified Eagle’s medium containing 10% fetal calf serum and 1% penicillin/streptomycin. One day before transfection the cells were plated on 35 mm fluorodishes (World Precision Instruments) coated with 100 µg/mL poly-D-lysine (Sigma Aldrich) and 4 µg/mL mouse laminin (Thermo Fisher Scientific). Cells were transfected with 120 ng PEX3-mEMerald-FKBP and BicD2NS-V5-FRB constructs per well as well as 20 pmol of either ON-TARGETplus Non-targeting siRNA #1 (Dharmacon) or SMARTpool: ON-TARGETplus PAFAH1B1 siRNA (Dharmacon) using Lipofectamine 2000 (Thermo Fisher Scientific).

Cells were labelled with Halo-JF549 (Janelia) and imaged after 48 hours using a 100x Apo TIRF NA 1.49 objective on a Nikon Ti2 microscope with a Yokogawa-X1 spinning disk confocal system, MLC400B laser engine (Agilent), Prime 95B back-thinned sCMOS camera (Teledyne Photometrics), piezo Z-stage (Mad City Labs) and stage top environmental chamber (Tokai Hit). Cells were screened for the presence of JF549 signal with the 560 laser line and then mEmerald was imaged at 2 frames per second, 100 ms exposure with the 488 laser line. Dimerization of FKBP-FRB was induced with 1 µM rapalog (Takara Bio). Images were analyzed in ImageJ. Kymographs were generated from 6-10 peroxisomes that moved directionally for >3 frames in each cell and velocity was calculated from kymographs using ImageJ macros as described^54^. Data plotting and statistical analyses were performed in Prism7 (GraphPad).

## References

1. Raaijmakers, J. A. & Medema, R. H. Function and regulation of dynein in mitotic chromosome segregation. Chromosoma 123, 407–422 (2014).

2. Reck-Peterson, S. L., Redwine, W. B., Vale, R. D. & Carter, A. P. The cytoplasmic dynein transport machinery and its many cargoes. Nature Reviews Molecular Cell Biology 19, 1–17 (2018).

3. Schlager, M. A., Hoang, H. T., Urnavicius, L., Bullock, S. L. & Carter, A. P. In vitro reconstitution of a highly processive recombinant human dynein complex. EMBO J 33, 1855–1868 (2014).

4. Mckenney, R. J., Huynh, W., Tanenbaum, M. E., Bhabha, G. & Vale, R. D. Activation of cytoplasmic dynein motility by dynactin-cargo adapter complexes. Science 345, 337–341 (2014).

5. Urnavicius, L. et al. Cryo-EM shows how dynactin recruits two dyneins for faster movement. Nature 554, 202–206 (2018).

6. Grotjahn, D. A. et al. Cryo-electron tomography reveals that dynactin recruits a team of dyneins for processive motility. Nat Struct Mol Biol 25, 203–207 (2018).

7. DeSantis, M. E. et al. Lis1 Has Two Opposing Modes of Regulating Cytoplasmic Dynein. Cell 170, 1197–1208.e12 (2017).

8. Lipka, J., Kuijpers, M., Jaworski, J. & Hoogenraad, C. C. Mutations in cytoplasmic dynein and its regulators cause malformations of cortical development and neurodegenerative diseases. Biochem Soc Trans 41, 1605–1612 (2013).

9. Olenick, M. A. & Holzbaur, E. L. F. Dynein activators and adaptors at a glance. J Cell Sci 132, jcs227132 (2019).

10. Reck-Peterson, S. L. et al. Single-molecule analysis of dynein processivity and stepping behavior. Cell 126, 335–348 (2006).

11. Trokter, M., Mücke, N. & Surrey, T. Reconstitution of the human cytoplasmic dynein complex. Proc Natl Acad Sci USA 109, 20895–20900 (2012).

12. Zhang, K. et al. Cryo-EM Reveals How Human Cytoplasmic Dynein Is Auto-inhibited and Activated. Cell 169, 1303–1314.e18 (2017).

13. Torisawa, T. et al. Autoinhibition and cooperative activation mechanisms of cytoplasmic dynein. Nat Cell Biol 16, 1118–1124 (2014).

14. Geiser, J. R. et al. Saccharomyces cerevisiae genes required in the absence of the CIN8-encoded spindle motor act in functionally diverse mitotic pathways. Mol Biol Cell 8, 1035–1050 (1997).

15. Xiang, X., Osmani, A. H., Osmani, S. A., Xin, M. & Morris, N. R. NudF, a nuclear migration gene in Aspergillus nidulans, is similar to the human LIS-1 gene required for neuronal migration. Mol Biol Cell 6, 297–310 (1995).

16. Liu, Z., Xie, T. & Steward, R. Lis1, the Drosophila homolog of a human lissencephaly disease gene, is required for germline cell division and oocyte differentiation. Development 126, 4477–4488 (1999).

17. Lam, C., Vergnolle, M. A. S., Thorpe, L., Woodman, P. G. & Allan, V. J. Functional interplay between LIS1, NDE1 and NDEL1 in dynein-dependent organelle positioning. J Cell Sci 123, 202–212 (2010).

18. Splinter, D. et al. BICD2, dynactin, and LIS1 cooperate in regulating dynein recruitment to cellular structures. Mol Biol Cell 23, 4226–4241 (2012).

19. Lenz, J. H., Schuchardt, I., Straube, A. & Steinberg, G. A dynein loading zone for retrograde endosome motility at microtubule plus-ends. EMBO J 25, 2275–2286 (2006).

20. Moughamian, A. J., Osborn, G. E., Lazarus, J. E., Maday, S. & Holzbaur, E. L. F. Ordered Recruitment of Dynactin to the Microtubule Plus-End is Required for Efficient Initiation of Retrograde Axonal Transport. Journal of Neuroscience 33, 13190–13203 (2013).

21. Lee, W. L. The role of the lissencephaly protein Pac1 during nuclear migration in budding yeast. J Cell Biol 160, 355–364 (2003).

22. Tsai, J.-W., Chen, Y., Kriegstein, A. R. & Vallee, R. B. LIS1 RNA interference blocks neural stem cell division, morphogenesis, and motility at multiple stages. J Cell Biol 170, 935–945 (2005).

23. Tanaka, T. et al. Lis1 and doublecortin function with dynein to mediate coupling of the nucleus to the centrosome in neuronal migration. J Cell Biol 165, 709–721 (2004).

24. Dix, C. I. et al. Lissencephaly-1 promotes the recruitment of dynein and dynactin to transported mRNAs. J Cell Biol 202, 479–494 (2013).

25. Reiner, O. et al. Isolation of a Miller-Dieker lissencephaly gene containing G protein beta-subunit-like repeats. Nature 364, 717–721 (1993).

26. Kim, M. H. et al. The structure of the N-terminal domain of the product of the lissencephaly gene Lis1 and its functional implications. Structure 12, 987–998 (2004).

27. Tarricone, C. et al. Coupling PAF signaling to dynein regulation: structure of LIS1 in complex with PAF-acetylhydrolase. Neuron 44, 809–821 (2004).

28. Toropova, K. et al. Lis1 regulates dynein by sterically blocking its mechanochemical cycle. Elife 3, 279 (2014).

29. Huang, J. J., Roberts, A. J. A., Leschziner, A. E. A. & Reck-Peterson, S. L. S. 6Lis1 acts as a ‘clutch’ between the ATPase and microtubule-binding domains of the dynein motor. 150, 975–986 (2012).

30. Yamada, M. et al. LIS1 and NDEL1 coordinate the plus-end-directed transport of cytoplasmic dynein. EMBO J 27, 2471–2483 (2008).

31. Mckenney, R. J., Vershinin, M., Kunwar, A., Vallee, R. B. & Gross, S. P. LIS1 and NudE induce a persistent dynein force-producing state. 141, 304–314 (2010).

32. Baumbach, J. et al. Lissencephaly-1 is a context-dependent regulator of the human dynein complex. Elife 6, 711 (2017).

33. Gutierrez, P. A., Ackermann, B. E., Vershinin, M. & Mckenney, R. J. Differential effects of the dynein-regulatory factor Lissencephaly-1 on processive dynein-dynactin motility. J Biol Chem 292, 12245–12255 (2017).

34. Redwine, W. B. et al. The human cytoplasmic dynein interactome reveals novel activators of motility. Elife 6, 379 (2017).

35. Hoogenraad, C. C. et al. Mammalian Golgi-associated Bicaudal-D2 functions in the dynein-dynactin pathway by interacting with these complexes. EMBO J 20, 4041–4054 (2001).

36. Hoogenraad, C. C. et al. Bicaudal D induces selective dynein-mediated microtubule minus end-directed transport. EMBO J 22, 6004–6015 (2003).

37. Kapitein, L. C. et al. Probing Intracellular Motor Protein Activity Using an Inducible Cargo Trafficking Assay. Biophys J 99, 2143–2152 (2010).

38. Tai, C.-Y., Dujardin, D. L., Faulkner, N. E. & Vallee, R. B. Role of dynein, dynactin, and CLIP-170 interactions in LIS1 kinetochore function. J Cell Biol 156, 959–968 (2002).

39. Sasaki, S. et al. A LIS1/NUDEL/cytoplasmic dynein heavy chain complex in the developing and adult nervous system. Neuron 28, 681–696 (2000).

40. Schmidt, H., Zalyte, R., Urnavicius, L. & Carter, A. P. Structure of human cytoplasmic dynein-2 primed for its power stroke. Nature 1–16 (2014). doi:10.1038/nature14023

41. Wang, S. et al. Nudel/NudE and Lis1 promote dynein and dynactin interaction in the context of spindle morphogenesis. Mol Biol Cell 24, 3522–3533 (2013).

42. Jha, R., Roostalu, J., Cade, N. I., Trokter, M. & Surrey, T. Combinatorial regulation of the balance between dynein microtubule end accumulation and initiation of directed motility. EMBO J 36, 3387–3404 (2017).

43. Egan, M. J. & Tan, K. Lis1 is an initiation factor for dynein-driven organelle transport. J Cell Biol 197, 971–982 (2012).

44. Zhang, J., Qiu, R., Arst, H. N., Penalva, M. A. & Xiang, X. HookA is a novel dynein-early endosome linker critical for cargo movement in vivo. J Cell Biol 204, 1009–1026 (2014).

45. Bielska, E. et al. Hook is an adapter that coordinates kinesin-3 and dynein cargo attachment on early endosomes. J Cell Biol 204, 989–1007 (2014).

46. Sheeman, B. et al. Determinants of S. cerevisiae dynein localization and activation: implications for the mechanism of spindle positioning. Curr Biol 13, 364–372 (2003).

47. Lee, W.-L., Kaiser, M. A. & Cooper, J. A. The offloading model for dynein function: differential function of motor subunits. J Cell Biol 168, 201–207 (2005).

48. Farkasovsky, M. & Küntzel, H. Yeast Num1p associates with the mother cell cortex during S/G2 phase and affects microtubular functions. J Cell Biol 131, 1003–1014 (1995).

49. Tang, X., Germain, B. S. & Lee, W.-L. A novel patch assembly domain in Num1 mediates dynein anchoring at the cortex during spindle positioning. J Cell Biol 196, 743–756 (2012).

50. Lammers, L. G. & Markus, S. M. The dynein cortical anchor Num1 activates dynein motility by relieving Pac1/LIS1-mediated inhibition. J Cell Biol 211, 309–322 (2015).

51. Markus, S. M. et al. Quantitative analysis of Pac1/LIS1-mediated dynein targeting: Implications for regulation of dynein activity in budding yeast. Cytoskeleton (Hoboken) 68, 157–174 (2011).

52. Wang, Y. et al. CRACR2a is a calcium-activated dynein adaptor protein that regulates endocytic traffic. J Cell Biol jcb.201806097 (2019). doi:10.1083/jcb.201806097

## Additional Methods References

53. Case, R. B., Pierce, D. W., Hom-Booher, N., Hart, C. L. & Vale, R. D. The directional preference of kinesin motors is specified by an element outside of the motor catalytic domain. Cell 90, 959–966 (1997).

54. Roberts, A. J., Goodman, B. S. & Reck-Peterson, S. L. Reconstitution of dynein transport to the microtubule plus end by kinesin. Elife 3, e02641 (2014).

55. Scheres, S. H. W. RELION: implementation of a Bayesian approach to cryo-EM structure determination. J Struct Biol 180, 519–530 (2012).

56. Lander, G. C. et al. Appion: an integrated, database-driven pipeline to facilitate EM image processing. J Struct Biol 166, 95–102 (2009).

57. Zheng, S. Q. et al. MotionCor2: anisotropic correction of beam-induced motion for improved cryo-electron microscopy. Nat Meth 14, 331–332 (2017).

58. Rohou, A. & Grigorieff, N. CTFFIND4: Fast and accurate defocus estimation from electron micrographs. J Struct Biol 192, 216–221 (2015).

59. Voss, N. R., Yoshioka, C. K., Radermacher, M., Potter, C. S. & Carragher, B. DoG Picker and TiltPicker: software tools to facilitate particle selection in single particle electron microscopy. J Struct Biol 166, 205–213 (2009).

60. Zivanov, J. et al. New tools for automated high-resolution cryo-EM structure determination in RELION-3. Elife 7, 163 (2018).

61. Punjani, A., Rubinstein, J. L., Fleet, D. J. & Brubaker, M. A. cryoSPARC: algorithms for rapid unsupervised cryo-EM structure determination. Nat Meth 14, 290–296 (2017).

